# Immunomodulatory activity of *Momordica charantia L. (Cucurbitaceae)* leaf diethyl ether and methanol extracts on *Salmonella typhi* infected mice and LPS-induced phagocytic activities of macrophages and neutrophils

**DOI:** 10.1101/731034

**Authors:** Oumar Mahamat, Hakoueu N. Flora, Tume Christopher, Kamanyi Albert

## Abstract

Infections due to *salmonella* strains constitute one of the major health problems in humans, particularly in Africa. Use of traditional herbs has proven effective in reducing the incidence of infection in some high-risk groups. To assess the effects of *Momordica charantia* leaf extracts that influence blood infestation, *in vitro* study of the effect on macrophages and neutrophils, and treatment of mouse model of *S. typhi* infection was done. Methanol and diethyl ether extracts were concerned by this study. *In vitro* study was to assess the effects of extracts on phagocytosis and related intracellular killing mechanisms of macrophages were examined. Later, mobilization of leukocytes and production of antibodies against *S. typhi* were measured followed by quantitating cultures evaluation of the blood infestation of orally inoculated mice with *S. thyphi*. Ingestion or attachment of carbon particles, production of superoxide anion, nitric oxide and that of lysosomal acid phosphatase by macrophages and neutrophils were significantly increased by methanol and diethyl extracts at concentrations ranging from 40 μg/ml to 640 μg/ml. Antibody titer and mobilization of leukocytes, particularly lymphocytes against *S. typhi* were highly increased by both methanol and diethyl extracts at concentrations of 500 and 1000 mg/kg. In the same the extracts have reduced the rate of blood infestation in mice inoculated with 10^8^ CFU of *S. typhi* for 28 days. Reduction in blood infestation rates was similar for levamisole mice group. Results of this study should prove useful of leave of *Momordica charantia* for treatment of infections by *salmonella* strains and for assessment of drugs for therapeutic intervention.

## Background

The immune system has a fundamental role in protecting the body against pathogenic microbial agents ^[1]^. Once activated, the immune system produces immediate response by the activation of immune component cells and production of various cytokines, chemokines and inflammatory mediators. In several conditions, the system is a target of a numerous drugs and herbs known as immunomodulators act by achieving immunostimulation (as in the treatment of AIDS) or achieving immunosuppression (e.g. the treatment of autoimmune disease) ^[2]^.

*Salmonella* infections are extremely common in the Cameroon. Frequently asymptomatic, salmonellosis imposes costs upon the public sector, on industry, in particular the wholesale and retail food industry, and very importantly upon the infected person and their family. Given both the wide distribution of *Salmonella* in foodstuffs and the frequency of asymptomatic *Salmonella* carriage, it is difficult to envision how any restaurant might prevent the occasional case of *Salmonella* transmission despite emphasis on hygienic practices. *Salmonella* infection is therefore a risk of everyday life, especially for persons who dine out frequently. As in all diseases, containment of *Salmonella* infection depends on an intact T-lymphocyte system including macrophage function. Persons with impaired T-cell function because of lymphoproliferative disorders, or immunosuppressive medication and persons with disorders that cause “macrophage blockade” such as hemoglobinopathies, malaria, schistosomiasis are well known to be persons at risk of serious consequences of *Salmonella* infection.

Modulation of immune response to alleviate disease conditions has long been of interest and increasingly recognized as a key component of effective disease control. Plant extracts have been widely investigated in the recent time in different parts of the world for their possible immunomodulatory properties ^[3,4]^. They are very helpful in prevention of infectious diseases, or acquired immunodeficiency ^[5]^.

Since most of the drugs currently available for treatment of salmonellosis are toxic, costly and no longer effective, attempts are being made in laboratories around the world to discover new, safer, more cost effective and more potent molecules from medicinal plants with an ethnomedical history. Many plant extracts with immunomodulatory activities can be of great help in the control of bacterial infection notably salmonellosis. Plants such as *Caesalpinia bonducella Flem* (*Caesalpiniaceae*), *Rhododendron spiciferum Franch (Ericaceae), Curcuma longa Linn (Zingiberaceae), Azadiracta indica A., Juss (Meliaceae), Boerhaavia diffussa Linn* (*Nyctaginaceae*) and *Ocimum sanctum Linn* (*Lamiaceae*) among others, are known to possess immunomodulatory activity ^[6]^.

*Momordica charantia* L. (*M. charantia*), is known to have both immunosuppressive and immunostimulant activities ^[7]^. Plant fruits were demonstrated to promote the phagocytic activity, activation of splenocytes ^[8–12]^. Several bioactive compounds of *M. charantia* fruit have been recorded in the literature. They are classified as carbohydrates, proteins, lipids and more ^[13–15]^. *M. charantia* contains triterpenoids ^[16–19]^, saponins ^[20–22]^, polypeptides ^[23]^, flavonoids ^[24]^, alkaloids ^[23]^ and sterols ^[18]^. Leave of *M. charantia* are used in Cameroonian traditional medicine to treat typhoid. But, the biological activities and mode of action of the plant extracts are poorly understood and may act directly or indirectly.

This work was therefore designed to study the immunomodulatory activity of methanol and diethyl ether extracts of *M. charantia* leave on *Salmonella typhi* infected mice and phagocytic cells with the aim of having a better understanding of the therapeutic of *M. charantia* against *salmonella* strains.

## Materials and methods

### Reagents and chemical

Various reagents and chemicals were used to prepare the extracts and for the assays. They include 3- (4,5-dimetilthiazol-2-yl) −2.5-diphenyl tetrazolium bromide (MTT), Roswell Park Memorial Institute (RPMI) medium, fetal bovine serum (FBS), para-nitrophenylphosphate (*P*-NPP), phosphate buffered saline (PBS), lipopolysaccharide (LPS), penicillin-streptomycin (Pen-Strep), neutral red (NR), sulfanilamide, naphthylethylenediamine dihydrochloride, dimethyl sulfoxide (DMSO), triton-100 and nitroblue tetrazolium (NBT) whose were purchased from Sigma Chemical, Germany. Methanol and diethyl ether used as solvents were obtained from Merck.

### Plant material

Leave of *M. charantia* were collected in May 2018 from Mbui Division, North West region, Cameroon. It was identified by Dr. TACHAM Walter, a botanist at Department of Biological Sciences, University of Bamenda, Cameroon. The identification was authenticated by the national herbarium in comparison with the collected material of Letouzey R6428, where the voucher specimen is registered under the following number: N°: 8095/REF/CAM.

### Experimental animals

Adult male out-bred albino mice (10-12 weeks old; 18–25 g) were used for the study. They were obtained from National Veterinary Laboratory, Garoua, Cameroon where they were raised under constant temperature (25-27°C) and light (12 hours light/dark). The animals were taken to the animal house of the Department of Biological Sciences, where they were given standard rodent feed and water ad libitum.

### Salmonella strain

*Salmonella typhi* was used for this study. It was isolated from clinically sick patients. It was maintained in the Department by serial cultures in SDS medium. This organism was grown on MacConkey’s agar, containing 2% agar, at 37°C. For 18 hours and harvested into sterile saline. The bacteria were washed three times in saline and finally suspended in 0.5% formalinized saline. This suspension was incubated at 37°C to kill the organisms and then tested for sterility. This sterile suspension constituted the stock antigen and was stored at 2°C. The antigens for inoculation into mice or for the agglutination test were diluted from the stock antigen with sterile saline to a density of tube of McFarlands nephelometer (10^9^ organisms per ml).

### Preparation of plant extracts

Fresh leave of *M. charantia* were washed with distilled water and dried at 30 °C. The dried leave were ground and weighed. Subsequently, the dried powder was extracted with 98% diethyl ether (ratio: 1:5) for 3 days at room temperature. The solvent-containing extract was then filtered and the filtrate was evaporated using a rotary evaporator to provide the diethyl ether extract (D-Extract). The residue was dried and extracted with 80% methanol with a ratio of 1:6. The filtrate was evaporated to provide the methanol extract (M-Extract). To further ensure that all the water was removed, the extracts were freeze dried using a dry ovum. The extract solutions were then prepared by dissolving 5 mg of extracts in the 0.25 ml DMSO and diluting with PBS to be used *in vitro* and *in vivo*.

### Ethical consideration and blood collection

Blood samples were collected from mice for experiment by cardiac puncture under anesthesia by mixture of Ketamine/Xylazine administered at 2 different doses (50 mg/kg–5 mg/kg), 0.1 ml/100g b.w. intraperitoneally. Animal studies were in compliance with the ethical procedures of the Animal Use and Care Committee, Faculty of Sciences, University of Bamenda, which corresponds with National Institutes of Health (NIH) guidelines ^[26]^.

### Cell preparation

Total white blood cells were obtained by collecting the plasma from heparinized mice blood and diluted in equal volume of RPMI-1640 medium. Cells were harvested after 2 successive washings by centrifugation (1800 rpm, 10 min) in medium. Peritoneal cells (neutrophils and macrophages) using elicitation methods after an intraperitoneally injection of fetal bovine serum ^[27]^. Neutrophils (PNs) were isolated 21 hours after injection of FBS while macrophages were harvested 3 days after injection of FBS. Ten millilitres of cold RPMI were injected in the peritoneal and the exudate was collected in sterile assay tube by syringe. The exudates containing the cells were then centrifuged at 1200 rpm for 10 min at 4°C, and the cells were washed twice and re-suspended in complete RPMI medium. Cells were counted using a haemocytometer, and viability was assessed by trypan blue exclusion. Cell number was therefore adjusted to the needed density.

### Preparation of Carbon Particle Suspension

A stable suspension of carbon particles was obtained by suspending the ultra-fine carbon powder in complete RMPI 1640. Then, the mass concentration of carbon particles in suspension was quantified by measuring the optical density at 800 nm with a spectrophotometer. A linear relationship has shown between mass concentration of carbon particles and optical density (Fig. 1).

**Fig. 1.**
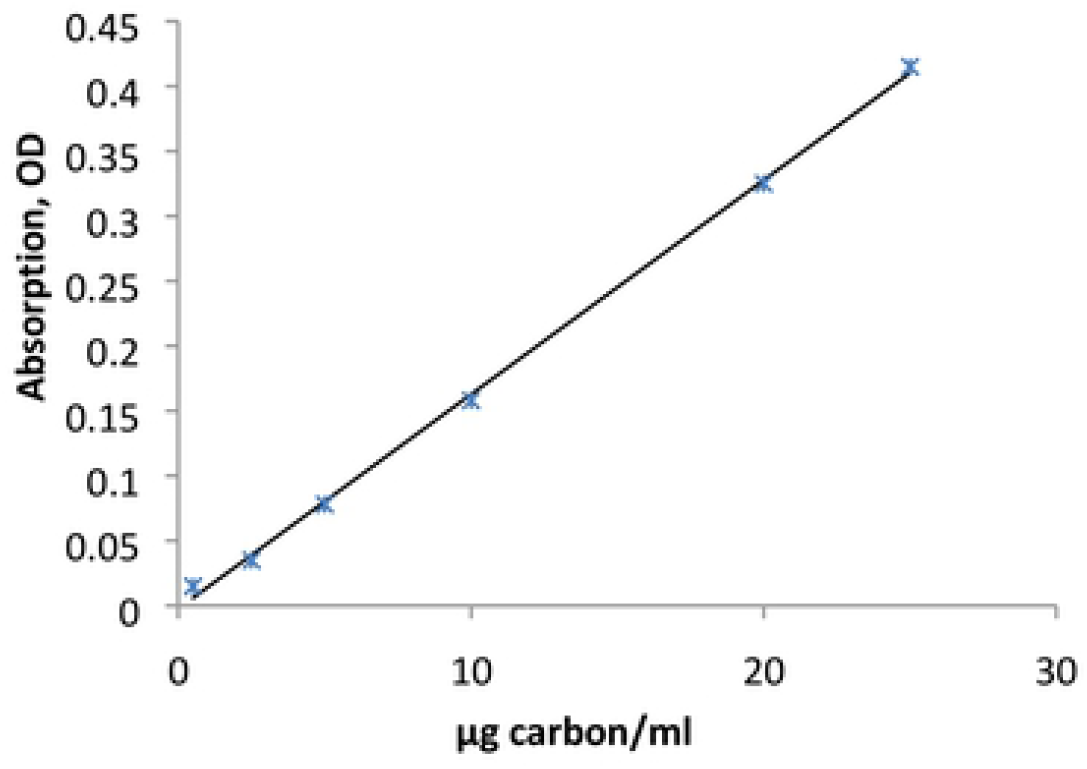
Correlation between the concentration of suspended carbon particles and optical density at 800 nm.

## In vitro immunomodulation studies

### Cell Stimulation

Neutrophils or macrophages were cultured with the extracts in 96-wells plate for then incubated at 37°C in a 5% CO_2_ humidified atmosphere in RMPI 1640 medium with the extracts for final concentrations (20 to 640 μg/ml) and LPS (4 μg/ml). Cell cultured added into well containing LPS only and medium only were taken as positive and negative controls, respectively. All solutions used were ensured to be lipopolysaccharide-free, and all assays were performed in triplicate and under sterile conditions. The cells were stimulated as described for all the various assays.

### Phagocytosis Assay

In order to examine whether extracts affect the phagocytic activity of macrophages and neutrophils, c*arbon Particles* were used as test particles in the studies of phagocytosis. A modification of the method as described by Margot ^[28]^ was used for the assessment of phagocytic function. Briefly, 1 ml of carbon particles in complete medium (25μg/ml) containing the extracts was added to test tubes with and without 1.5 x 10^6^ peritoneal cells. The samples were placed in a shaking water bath at 37°C for 12 h in order to obtain cells (macrophages and neutrophils) with attached and ingested carbon particles. The tubes were then centrifuged at 1000 tr/min for 15 min. The supernatants from the tubes with and without cells were measured in the spectrophotometer and the difference in carbon particle concentration was taken as a measure of attached and ingested particles.

### Assay of Oxidative Metabolism

The oxidative metabolism was measured by using the ability of the produced superoxide to reduce yellow nitroblue tetrazolium (NBT) to blue formazan. The assay was done to scrutinize the production of superoxide anion which is proportionally to reduction of the NBT. The assay was performed as previously described ^[29]^ in macrophages and neutrophils (1.5 ×10^5^ cells/mL) from three rats. After 48 h of incubation with or without extracts at 37°C in a 5% CO_2_ humidified incubator, 50 μl of freshly prepared 1.5 mg/ml NBT dye solution was added. Then, the adherent phagocytes were rinsed vigorously after incubation for 60 min with RPMI medium and washed four times with methanol. After air drying, 2 M KOH and DMSO were added, and the absorbance was measured at 570 nm using a microplate reader.

### Nitric Oxide Determination

Nitric oxide (NO) concentration was determined after 48 hours incubation with samples in 96-well plates, NO levels in each well were identified using the Griess reagent according to a previous study ^[29]^. After pre-incubation of macrophages or neutrophils (1.5 ×10^5^ cells/mL) with LPS (4 μg/mL) for 48 h, the quantity of nitrite in the culture medium was measured as an indicator of NO production. Amounts of nitrite, a stable metabolite of NO, were measured using Griess reagent (1% sulfanilamide and 0.1% naphthylethylenediamine dihydrochloride in 2.5% phosphoric acid) in 100 μl of the supernatant. The supernatant was mixed with equal volume of Griess reagent and the absorbance at 540 nm was measured in a microplate reader after incubation for 30 min.

### Acid Phosphatase Determination

Macrophages or neutrophils (1.5 ×10^5^ cells/mL) were incubated with LPS (4 μg/mL). After the desired length of time (48 hours) of incubation, the culture media were removed. Plates were washed twice with PBS. The adherent cells were then lysed with 100 μl of cold lysis buffer (0.2% Triton X-100 in 0.05M acetate buffer pH 5.4) and sonication (the cell plates were placed on ice) for 30 minutes. Cell extracts (100 μL) were mixed with 100 μl of an assay mixture containing 20 μl glacial acetic acid, 6 mg/mL *P*-NPP, and 0.1 mol/L acetate buffer (pH, 5.4). At 65 minutes, the reaction was stopped by the addition of 100 μL 1 N NaOH. The color was measured at 410 nm in a microplate reader. The activity of the extract (% stimulation) was calculated using the absorbance of treated and untreated wells.

### Expression of Percentage of stimulation

The assay was carried out in triplicates. The activity of extract was expressed as percentage of stimulation in each of the test well. The % of stimulation was calculated according to the following formula:

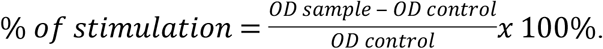

The OD control is the optical density of negative control and OD sample, the optical density of sample.

## In vivo immunomodulation studies

### In vivo leucocytes mobilization

Leucocytes mobilization method as described by Ribeiro ^[30]^ was used with few modifications to study the effect of the extracts on *in vivo* leucocytes migration induced by inflammatory stimulus. Thirty adult male mice infected intraperitoneally with *Salmonella typhi* were divided into five groups of 6 each. On day 3 and 7 post infection, three groups were given 250, 500 and 1000 mg/kg weight of extract, respectively by gavage. One of the remaining groups has received 7.5 mg/kg body weight of Levamisole and the last group was left as a control. One hour later, each mouse received intraperitoneal injection of 0.5 mL of 3% agar suspension in normal saline. Four hours later, the mice were sacrificed under anaesthesia and the peritoneum washed with 5 mL of phosphate buffer saline containing 0.5 mL of 10% EDTA. The peritoneal fluid was recovered and total leucocytes counts (TLC) determined with haemocytometer and the differential cell count was determined by microscopic counting of Giemsa stained perfusate smear on glass slide.

### Antibody Titrations

*Salmonella typhi* agglutinins were measured by an agglutination test using an antigen equivalent to 10^9^ organisms per ml and doubling dilutions of anti-sera starting as 1/10. The tests were incubated at 37°C for 12 hours, and then read. A second reading was made after a further incubation for 15 hours at room temperature. The titer of the serum was taken as the highest dilution in which definite agglutination was detected.

### Infection model

Inoculum contained a dose of *S. typhi* (10^8^ CFU in 1 ml of saline) was given to mice orally by gavage. Twenty seven days after inoculation, the animals were euthanized with diethyl ether and evaluated for blood infestation by *S. typhi*. Heparinized blood samples were obtained by tail vein puncture, and duplicate 100-μl aliquots were plated on SDS. The carriage rate was defined as the number of animals with ≥1 CFU divided by the total number of mice per group.

### Data Analysis

Experiments were done in triplicate or quadruplicate. Experimental results are presented as mean ± standard deviation. Data analysis was performed by one-way analysis of variance test followed by Tukey’s multiple comparison tests. Analysis was done using the program Graphpad Prism version 5.0. A P-value < 0.05 was considered statistically.

## Results

### Carbon Particles uptake

Table 1 shows a comparison of the percentage of carbon particles uptaken by macrophages and neutrophils treated with M-Extract and D-Extract carbon particles. In all samples, exposure to extracts increased ingested particles per peritoneal macrophages and neutrophils, the accumulated attachment, and the ingested fraction. The values for all extract concentrations (40, 160 and 640 μg/ml) were highly different compared to untreated macrophages and neutrophils (P<0.05) showing that PNs and PMs have reduced the particles put in presence. Respectively for PMs and PNs the reduced carbon particles was 1l.84 and 14.87μg/10^6^ cells at concentrations 640 μg/ml.

**Table. 1.** Effect of methanol and diethyl ether extracts of *M. charantia* on the uptake of carbon particles by macrophages and neutrophils.

| Samples | Methanol extract |  | Diethyl ether extract |  |
| --- | --- | --- | --- | --- |
|  | % reduction of OD | µg/10<sup>6</sup> cells | % reduction of OD | µg/10<sup>6</sup> cells |
| Attached and Ingested particles per Macrophages |  |  |  |  |
| Controls |  |  |  |  |
| (-) control | 31.87±1.54 | 5.31 | 31.87±1.54 | 5.31 |
| (+) control | 52.99±1.154*** | 8.83 | 52.99±1.154*** | 8.83 |
| Extracts (µg/ml) |  |  |  |  |
| 40 | <sup>a</sup>53.98±1.32*** | 8.99 | <sup>a</sup>42.01±1.94*** | 7.00 |
| 160 | <sup>b</sup>70.81±1.12*** | 11.80 | <sup>b</sup>57.25±1.66*** | 9.54 |
| 640 | <sup>c</sup>85.53±0.83*** | 14.25 | <sup>c</sup>71.07±0.41*** | 11.54 |
| Attached and Ingested particles per Neutrophils |  |  |  |  |
| Controls |  |  |  |  |
| (-) control | 43.14±3.70 | 7.19 | 43.14±3.70 | 7.19 |
| (+) control | 82.67±1.79*** | 13.77 | 82.67±1.79*** | 13.77 |
| Extracts (µg/ml) |  |  |  |  |
| 40 | <sup>a</sup>85.63±0.14*** | 14.27 | <sup>a</sup>83.08±0.97*** | 13.84 |
| 160 | <sup>ab</sup>87.51±0.92*** | 14.58 | <sup>ab</sup>86.12±0.09*** | 14.35 |
| 640 | <sup>b</sup>90.46±0.98*** | 15.07 | <sup>b</sup>89.23±0.56*** | 14.87 |

**Notes.** Three samples in three mice were studied. The amounts of carbon taken up by the cells were estimated by measurements of optical density (OD) of the carbon particle suspension (25 µg/ml) without the (1.5x10<sup>6</sup> cells/ml) were added and after the cells were removed by centrifugation. The asterisks indicate the significant difference relative to (-) control determined by Tukey's test ( $p \leq 0.05$ ). In the same column, the letters indicate the significant difference determined by Tukey's test ( $p \leq 0.05$ ).

### Superoxide anion production

*M. charantia* through D-Extract and M-Extract increased NBT dye reduction relative to untreated cells of LPS-induced mouse neutrophils in a dose dependent fashion at concentration ≥ 320 μg/ml (Fig. 2). In detail, in presence of D-Extract, the % of stimulation of NBT dye reduction in neutrophils at concentration of 640 and 320 μg/ml for 48 h was 68.48 ± 3.74%, 58.85 ± 3.23% and 46.52 ± 3.18% of LPS-control, respectively. With M-Extract at concentration of 640 and 320 μg/ml, the NBT dye reduction was 88.28±7.43%, 74.47 ± 6.72% and 50,00 ± 9,08% of LPS-control, respectively. A dose dependent increase of NBT dye reduction was also observed in macrophages. A significant increase was observed with D-Extract at 640 μg/ml where the % of stimulation was 45.83 ± 13.88% and 7.87 ± 2.89% for LPS-control. While, the incubation with M-Extract has resulted in augmentation of NBT reduction at 640, 320 and 150 μg/ml where the effect were 31.11 ± 2.77%, 21.01 ± 2.12%, 10.09 ± 0.80% and 6.01 ± 0.80% for LPS-control, respectively.

**Fig. 2.**
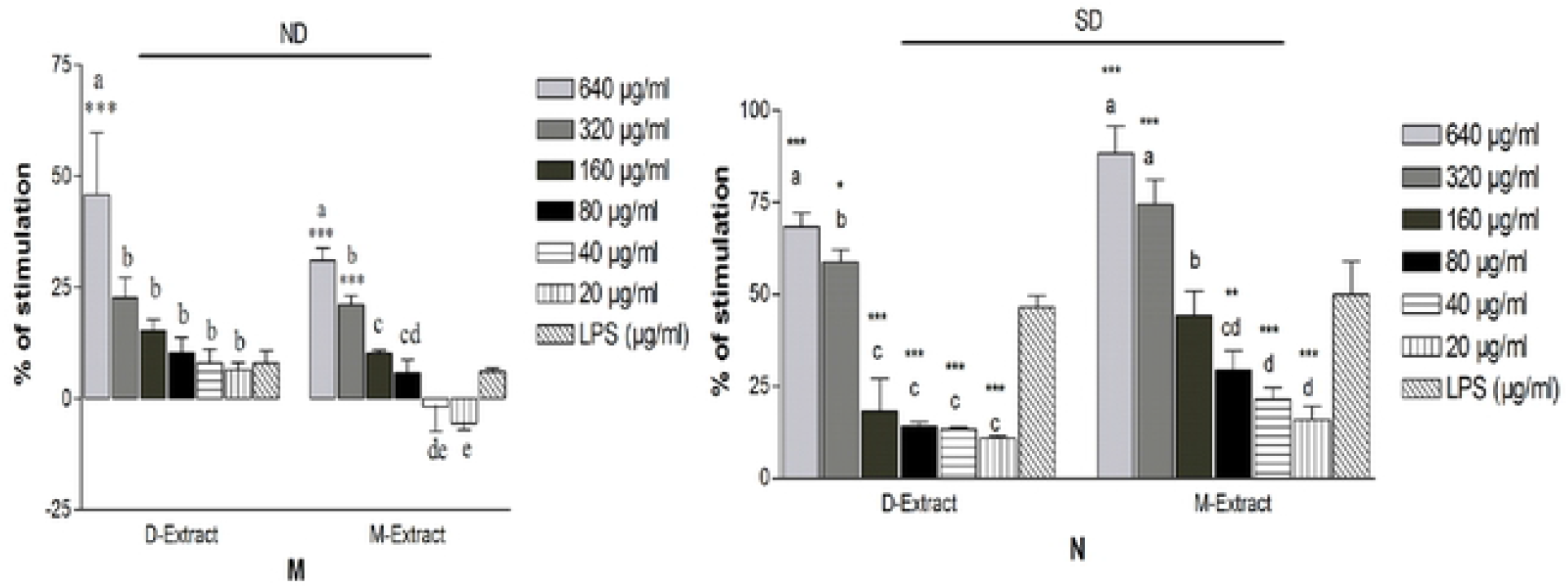
Stimulatory properties of the diethyl ether extracts (D-extract) and methanol extract (M-extract) on superoxide anion production by lipopolysaccharide (LPS)-induced peritoneal mouse macrophages (M) and neutrophils (N). The histogram expressed the mean ± SD (n = 4). The asterisks indicate the significant difference relative to LPS-control (4μg/ml) determined by Tukey’s test (p≤0.05). The letters indicate the significant difference determined by Tukey’s test (p≤0.05). SD and ND indicate respectively the significant difference and the absence of difference in the action of the two extracts determined by Tukey’s test (p ≤ 0.05).

### Nitric oxide concentration

The results of this study as presented in figure 3 demonstrated that *M. charantia* through its D-extract and M-extract at various concentrations caused a significant increase of NO production by both neutrophils and macrophages when compared with the LPS control (Fig.4). For the M-extract which has stimulated the NO production in macrophages only, the detail of the production at concentrations of 640, 320, 150, 80, 40 and 20 μg/mL for 48 h were 232.98 ± 28.34%, 128.76 ± 5.52%, 94.27 ± 7.38%, 83.50 ± 6.35%, 47.97 ± 12.11%, 34.83 ± 18.56% and respectively, compared to 89.32 ±1.84% of the control group treated with media only. On neutrophils, both D-extract and M-extract has promoted the production of NO (p < 0.05). In detail, the production of NO in LPS-induced neutrophils incubated with D-extract at concentrations of 640, 320, 150, 80, 40 and 20 μg/mL for 48 h were 108.80 ± 1.40%, 84.70 ± 12.21%, 60.52 ± 20.21%, 23.63 ± 3.11%, 24.14 ± 8.70%, 4.67 ± 1.80% and 37.58 ± 3.89% of the group treated with LPS only, respectively. The production of NO in LPS-induced neutrophils incubated with M-extract at the same concentrations were 297.91 ± 62.21%, 171.45 ± 39.42%, 129.50 ± 27.07%, 79.37 ± 12.47%, 47.54 ± 14.51%, 36.03 ± 19.06% and 37.58 ± 3,89% of the group treated with LPS only, respectively.

**Figure 3:**
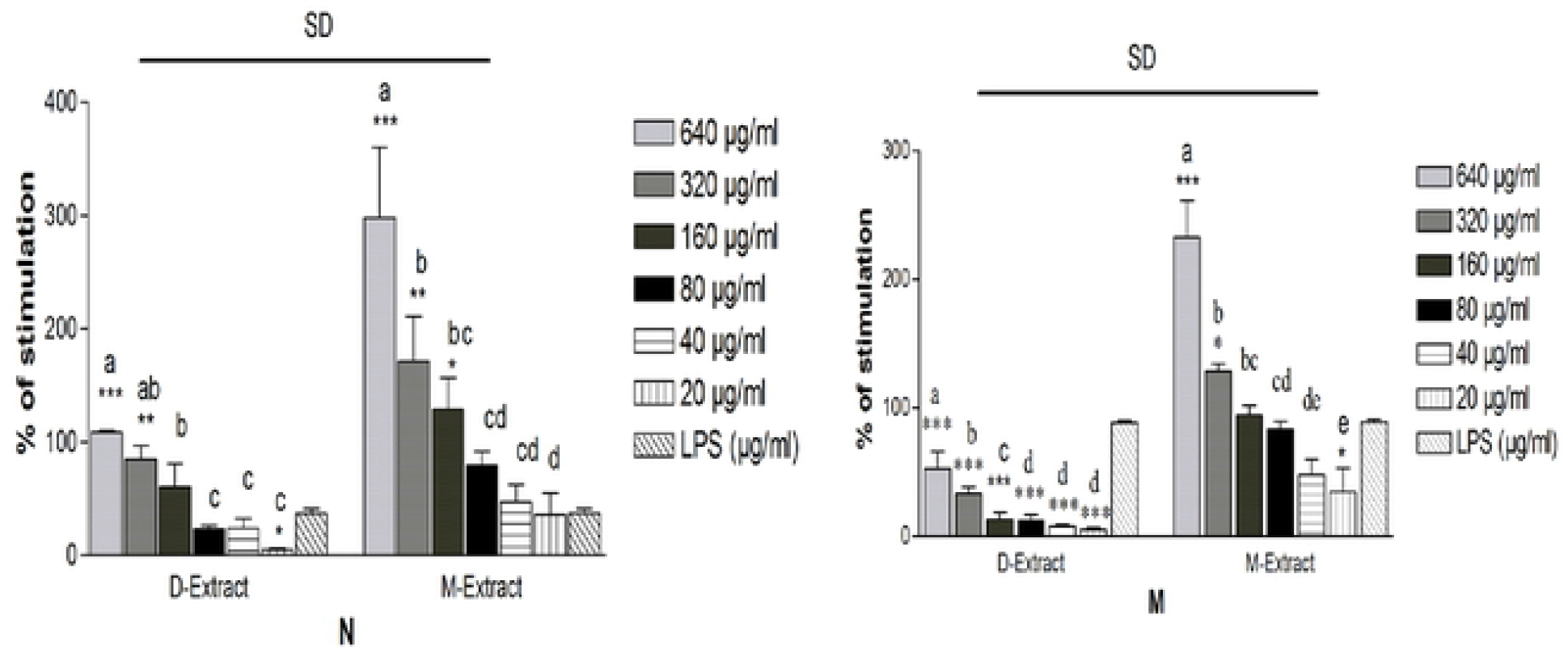
Stimulatory properties of the diethyl ether extracts (D-extract) and methanol extract (M-extract) on nitric oxide production by lipopolysaccharide (LPS)-induced peritoneal mouse macrophages (M) and neutrophils (N). The histogram expressed the mean ± SD (n = 4). The asterisks indicate the significant difference relative to LPS-control (4μg/ml) determined by Tukey’s test (p ≤ 0.05). The letters indicate the significant difference determined by Tukey’s test (p ≤ 0.05). SD indicates the significant difference in the action of the two extracts determined by Tukey’s test (p ≤ 0.05).

**Figure 4:**
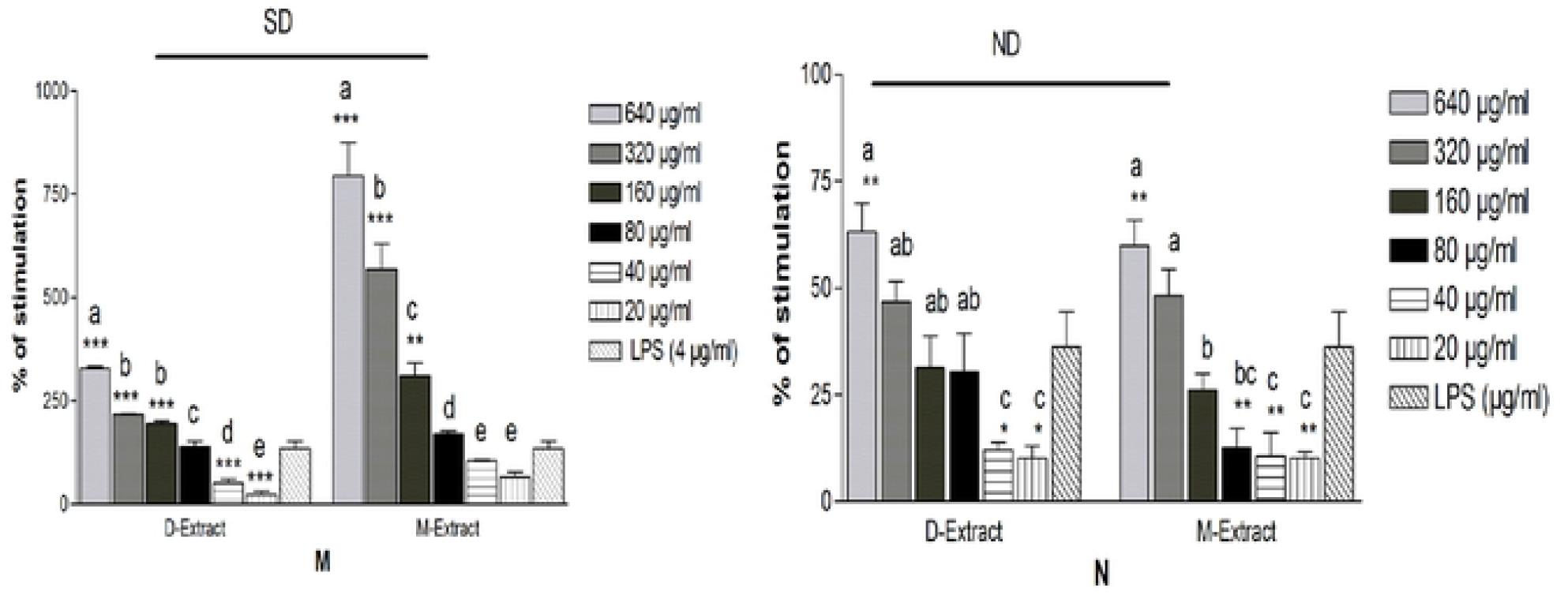
Stimulatory properties of the diethyl ether extracts (D-extract) and methanol extract (M-extract) on acid phosphatase activity in lipopolysaccharide (LPS)-induced peritoneal mouse macrophages (M) and neutrophils (N). The histogram expressed the mean ± SD (n = 4). The asterisks indicate the significant difference relative to LPS-control (4μg/ml) determined by Tukey’s test (p ≤ 0.05). The letters indicate the significant difference determined by Tukey’s test (p ≤ 0.05). SD and ND indicate respectively the significant difference and the absence of difference in the action of the two extracts determined by Tukey’s test (p ≤ 0.05).

### Acid phosphatase activity

Acid phosphatase activity was measured in LPS-induced neutrophils and macrophages after two days of incubation with the extracts. Our results showed that AcP activity, by with the presence of D-Extract and M-Extract was significantly increased in a dose-dependent manner (Fig. 4). The 640 μg/mL extracts only showed significantly higher augmentation of AcP activity in neutrophils (p < 0.05). At 640 μg / ml of D-Extract and M-Extract, the % of stimulation was 63.28 ± 6.53%, 59.90 ± 6.03% and 36.23 ± 8.19% for LPS-control, respectively. Both the diethyl ether and methanol extracts showed significant augmentation dose dependent of AcP activity in macrophages (p < 0.05). The percentage of stimulation in LPS-induced macrophages incubated with D-extract at concentrations of 640, 320 and 150 μg/mL were 328.99 ± 3.58%, 216.89 ± 2.15% and 194.64 ± 5.84% respectively. In cells incubated with M-Extract at concentrations of 640, 320 and 150 μg/mL the percentage of stimulation were 794.33 ± 79.26%, 568.19 ± 60.58%, 309.49 ± 32.46% and 134.15 ± 17.54% of the LPS-treated group value, respectively.

### Leucocytes mobilization

The effect of the extracts on in vivo leucocytes mobilization led to augmentation in total leucocytes count. With M-extract, the concentrations 500 mg/kg and 1000 mg/kg significantly increased the total leucocytes count from control group, with the effect higher than levamisole group. In differential leucocytes mobilization, all the tested doses significantly increased the lymphocyte count, while monocyte count has been increased in 1000 mg/kg group. In contrast, the basophil and eosinophil of the extract treated group showed decrease in number when compared with the control group (Table 2). Concentrations 500 mg/kg and 1000 mg/kg of D-extract also significantly increased the total leucocytes count in concentration dependent manner compared to control group. In differential leucocytes mobilization, all the tested doses significantly increased the lymphocyte count. while, the basophil and eosinophil showed decrease in number when compared with the control group (Table 3).

**Table 2.** Effect of *M. charantia* leaf methanol extract on total and differential leucocytes mobilization (cells/ml) in mice

|  | **Extract** |  |  | **Levanisole** | **Control** |
| --- | --- | --- | --- | --- | --- |
|  | **250 mg/kg** | **500 mg/kg** | **1000 mg/kg** | **7.5 mg/kg** | **-** |
| **TLC** | 1416,6±166,55a | 1675,6±144,60**b | 1981±461,18***b | 1184,4±149,24 | 962,28±144,41 |
| **NEUT** | 318,35±147,30<br>(22,09) | 391,04±167,52<br>(22,95) | 462,40±132,83<br>(23,44) | 283,69±75,77<br>(23,82) | 445,53±197,16<br>(45,68) |
| **LYMPH** | 1069,60±146,38**a<br>(75,88) | 1266,06±111,73***ab<br>(75,97) | 1467,92±364,80***b<br>(73,85) | 892,12±108,61*<br>(75,47) | 477,33±153,21<br>(50,186) |
| **BASO** | 4,38±4,51<br>(0,29) | 0,03±0,07**<br>(0,00) | 0.00±0.00**<br>(0.00) | 0,46±0,60**<br>(0,04) | 10,11±6,24<br>(1,07) |
| **MONO** | 15,10±5,12a<br>(1,07) | 1,22±1,14*b<br>(0,07) | 30,95±7,59***c<br>(1,61) | 1,64±0,94*<br>(0,13) | 14,06±7,90<br>(1,50) |
| **EOSIN** | 0,03±0,06<br>(0,03) | 4,92±2,44<br>(0,29) | 3,93±2,34<br>(0,21) | 5,38±7,09<br>(0,42) | 9,27±6,00<br>(0,93) |

Mean ± SD with different superscript letters are significantly different (p < 0.05).

**Table 3.**
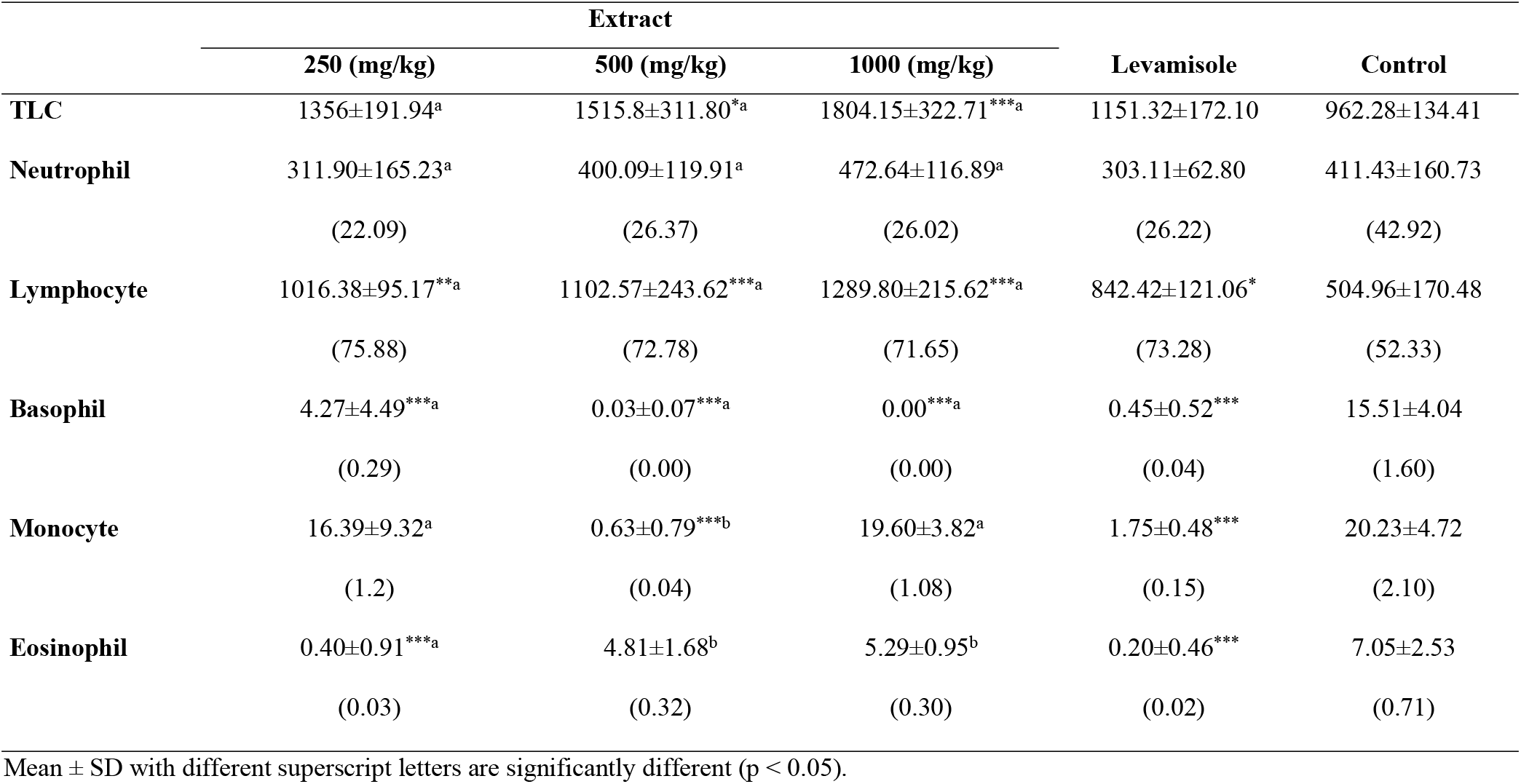
Effect of *M. charantia* leaf diethyl ether extract on total and differential leucocytes mobilization (cells/ml) in mice

|  | Extract |  |  | Levamisole | Control |
| --- | --- | --- | --- | --- | --- |
|  | 250 (mg/kg) | 500 (mg/kg) | 1000 (mg/kg) |  |  |
| **TLC** | 1356±191.94<sup>a</sup> | 1515.8±311.80<sup>*a</sup> | 1804.15±322.71<sup>***a</sup> | 1151.32±172.10 | 962.28±134.41 |
| **Neutrophil** | 311.90±165.23<sup>a</sup><br>(22.09) | 400.09±119.91<sup>a</sup><br>(26.37) | 472.64±116.89<sup>a</sup><br>(26.02) | 303.11±62.80<br>(26.22) | 411.43±160.73<br>(42.92) |
| **Lymphocyte** | 1016.38±95.17<sup>**a</sup><br>(75.88) | 1102.57±243.62<sup>***a</sup><br>(72.78) | 1289.80±215.62<sup>***a</sup><br>(71.65) | 842.42±121.06<sup>*</sup><br>(73.28) | 504.96±170.48<br>(52.33) |
| **Basophil** | 4.27±4.49<sup>***a</sup><br>(0.29) | 0.03±0.07<sup>***a</sup><br>(0.00) | 0.00<sup>***a</sup><br>(0.00) | 0.45±0.52<sup>***</sup><br>(0.04) | 15.51±4.04<br>(1.60) |
| **Monocyte** | 16.39±9.32<sup>a</sup><br>(1.2) | 0.63±0.79<sup>***b</sup><br>(0.04) | 19.60±3.82<sup>a</sup><br>(1.08) | 1.75±0.48<sup>***</sup><br>(0.15) | 20.23±4.72<br>(2.10) |
| **Eosinophil** | 0.40±0.91<sup>***a</sup><br>(0.03) | 4.81±1.68<sup>b</sup><br>(0.32) | 5.29±0.95<sup>b</sup><br>(0.30) | 0.20±0.46<sup>***</sup><br>(0.02) | 7.05±2.53<br>(0.71) |

Mean ± SD with different superscript letters are significantly different (p < 0.05).

### Anti-Salmonella typhi antibody titers

Antibodies against *Salmonella typhi* were determined by HA technique. The results showed that HA titers increased after treatment at day 14 and day 28 in animal. In the M-extract treated group, the primary antibody titers ranged from 160 to 640 (mean of 228 and 17 in the control group) at 500 mg/kg, whereas that of animal receiving 1000 mg/kg was from 320 to 1280 (mean of 1280). For this extract also an increase of antibodies at day 28 with a mean of 1664 and 4096 at the doses of 500 and 1000 mg/kg, whereas that of the control group was 44. An increase in antibody titers was also seen in levanisole treated group as well as at day 14 with a mean of 960 and 2304 at day 28 (Fig. 5).

**Fig. 5:**
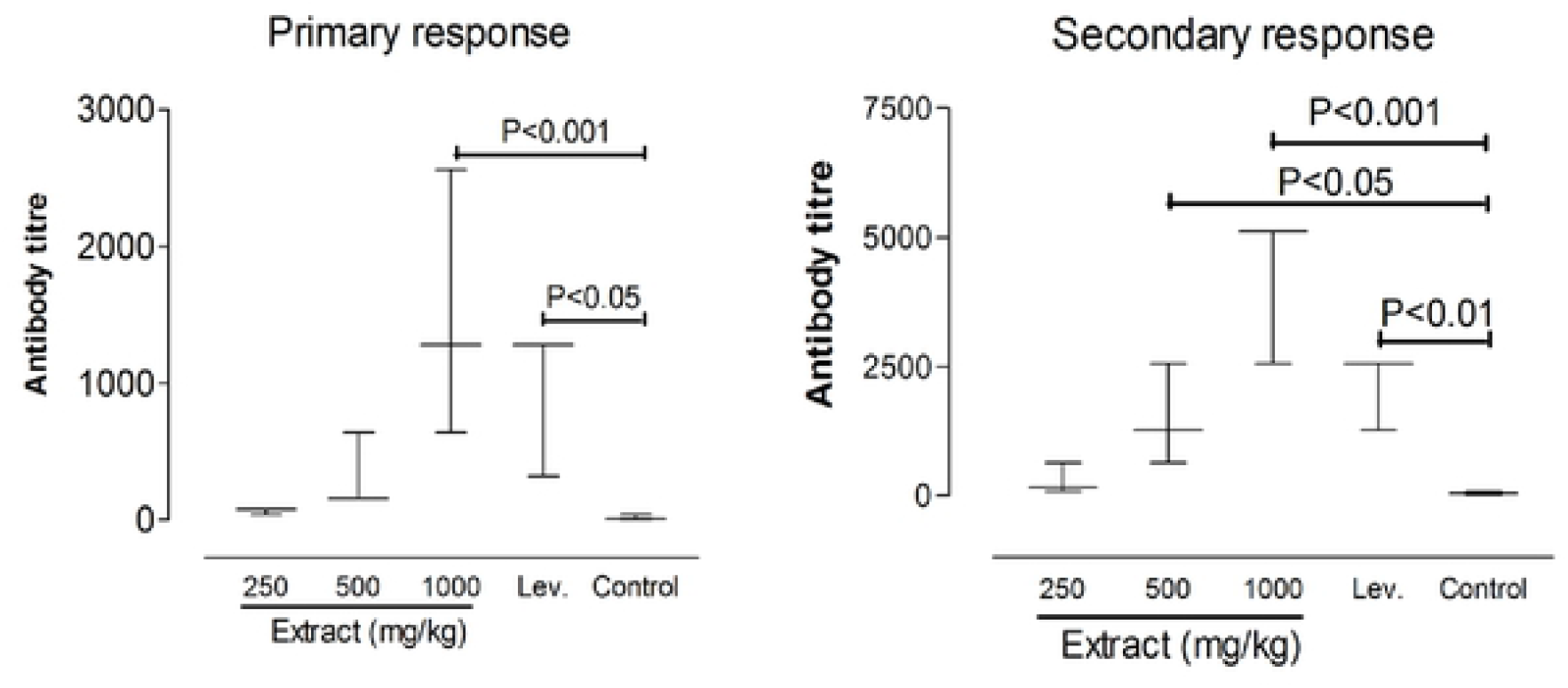
Effect of methanol extract on the antibody titre against *Salmonella typhi in mice*. The probability is the result of turkey test indicating the difference of the doses against control.

Besides, the results of this study in animals treated with D-extract, antibody titer against *S. typhi* was significantly increased in group receiving 1000 mg/kg with a mean of 168 at day 14. While at day 28, increase of antibody titer was seen in groups receiving the extract at 500 and 1000 mg/kg with means of 416 and 1152 respectively. HA titer means at day 14 and 28 were 18 and 448 in the levanisole treated group, whereas the means were 8 and 11 in the control group (Fig.6).

**Fig. 6:**
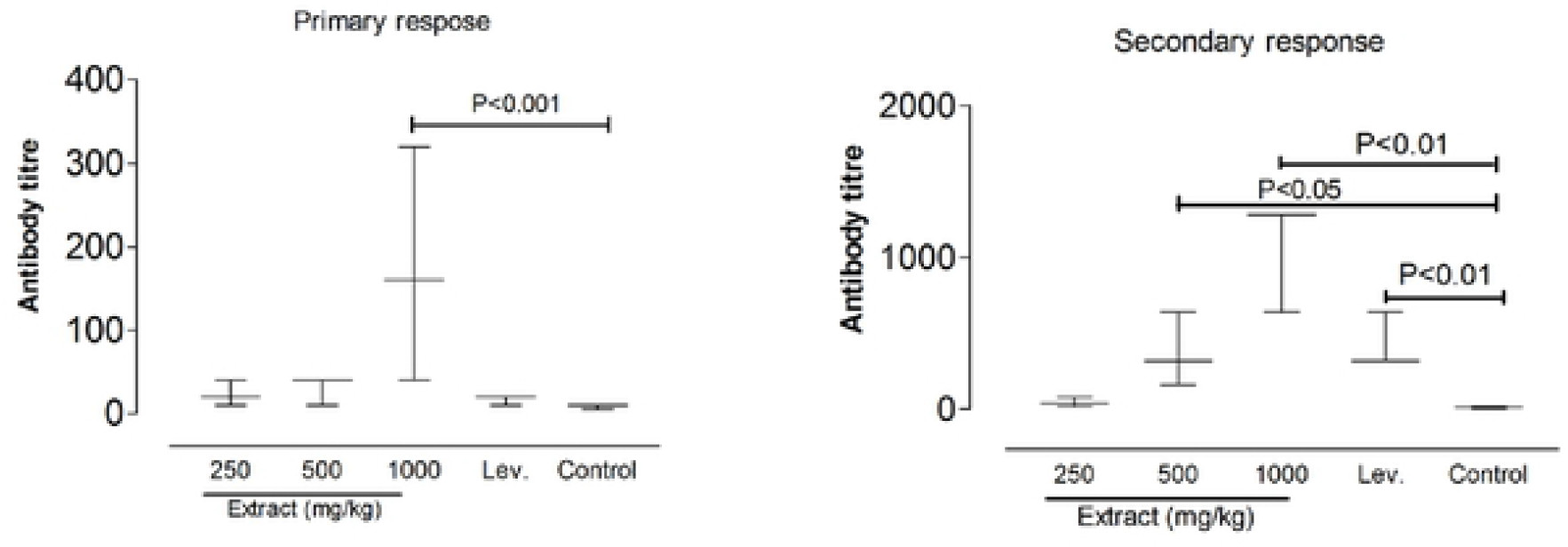
Effect of diethyl ether extract on the antibody titre against *Salmonella typhi in mice*. The probability is the result of turkey test indicating the difference of the doses against control.

### Effect on the blood infestation rate

At the time of blood collection after three experiment carry out independently, the percentage of mice having blood infested with ≥1 CFU of *S. typhi* in 500 and 1000 mg/kg treated groups was significantly lower than that of control groups. The proportion of infected animal was found to be 83.33% and 44.44% respectively for 500 and 1000 mg/g/kg against 94.44% for the control group. Levamisole also significantly decreased the proportion of mice with infested blood (38.88%) compared to control group (Fig. 7).

**Fig. 7:**
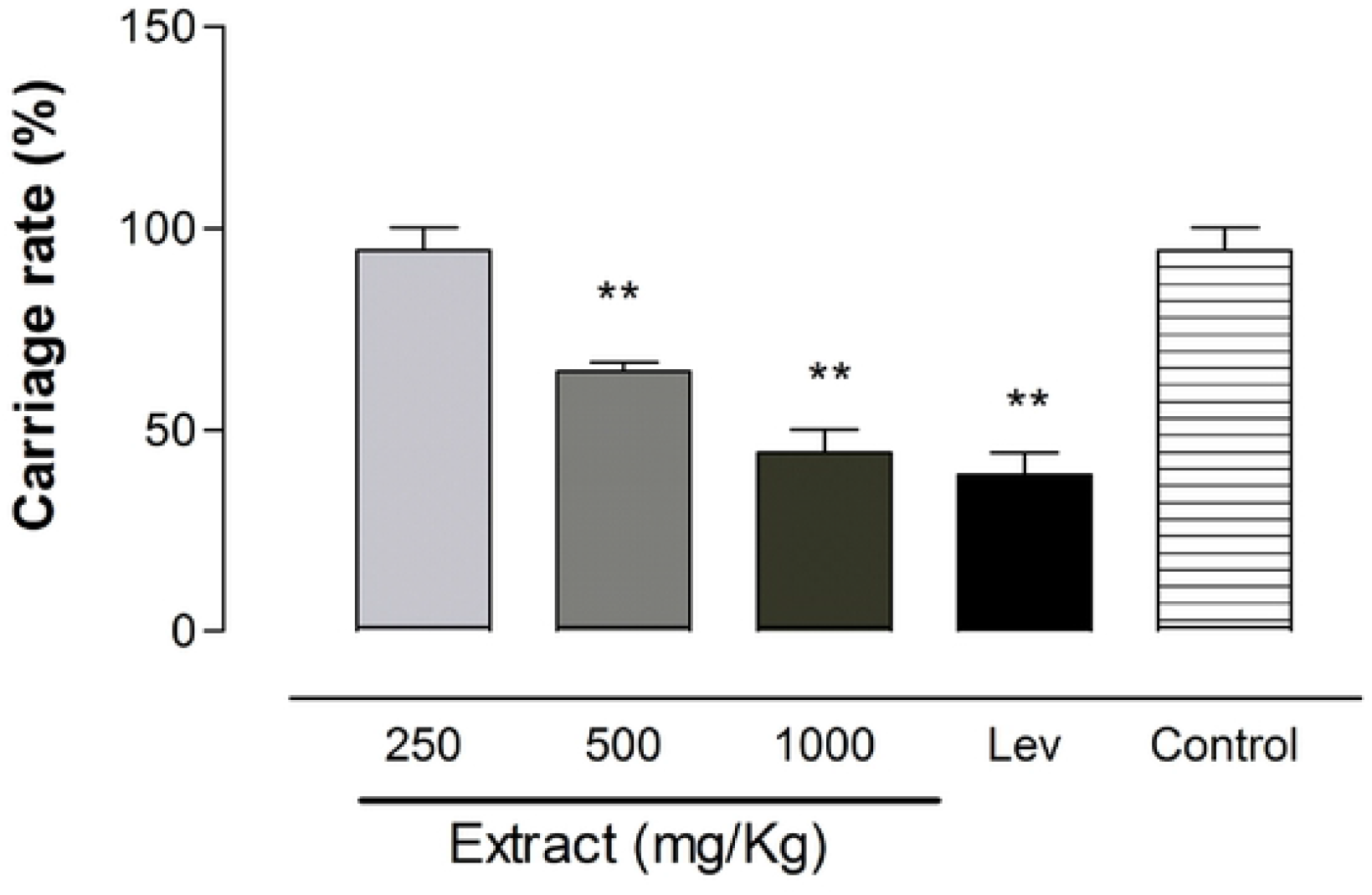
Blood infestation of mice with *Salmonella typhi* following the extract doses. The *P* values derived from the statistical comparison of carriage rate from the groups of untreated or treated extract-treated mice are shown below the graph. A P<0.05 was considered significant.

## Discussion

Many studies have addressed the immunomodulatory activities of *M. charantia* but, very little is known about phagocytic mechanisms and factors that influence blood infestation by bacterial strain. Passage of Salmonella strain in the blood is a multifactorial process that requires a variety of adaptive mechanisms, including adherence to host tissues, and host defenses. Identifying the immunotherapeutic factors of *M. charantia* that influence blood infestation by this bacterium is a best way for the use of tis plant. We have studied the phagocytic activity of the M-extract and D-extract of leaf of *M. charantia* in vitro in macrophages and neutrophils. Furthermore, we treated with extracts oral infected mice to conclude about the role *M. charantia* leaf in treatment of salmonellosis as pretending the traditional therapists in North West region of Cameroon.

It has been found that exposure of neutrophils and macrophages to M-Extract and D-Extract stimulates both their capacity to ingest foreign particles and their intracellular killing activities. This activity demonstrates an immunostimulation of phagocytosis ^[5,31]^. After exposure to extracts neutrophils and macrophages were found to be more functionally, as shown by the release of oxygen radicals. The results follow the earlier studies exhibiting *in vivo* and *in vitro* stimulatory effect of phagocytic activity by fruits of this plant ^[8–10]^. Production of the reactive oxygen species (ROS) is known to be increased during infection through activation of NADPH oxidase, therefore *M. charantia* leaf may be stimulated synthesis or activation of NADPH oxidase.

Furthermore, it has been demonstrated that the extracts stimulate the production of nitric oxide. The role of this nitrogen reactive specie is well known during immune reactions ^[32]^, and its production by neutrophils and macrophages is a result of iNOS synthesis. The effect of *M. charantia* leaf extracts in salmonellosis may be also attributed to that reactive specie production.

Under circumstances as bacteria activation, neutrophils or macrophages synthesis acid phosphatase, what it has been seen in neutrophils and macrophages exposed to extracts. Thus, *M. charantia* leaf might contribute in elimination of bacteria by stimulating lysosomal enzymes synthesis.

The result of this present study indicated that the extract increased the total leucocytes count of the perfusate when compared with the control group. Leucocytes migration is important for the transport of immunological information between different compartments of the immune system ^[33]^, suggesting that are stimulating the response to *S. typhi*. In addition, it has been found that all the tested doses improved the lymphocyte count. These cells are more important in the production of immunomodulatory cytokines and production of antibodies. Particularly, Th2 lymphocytes are direct leukocyte producing of IL-4, suggesting a prominent role for these cells of the adaptive immune system in the biology of B cells notably production of immunoglobulins what can be attributed to *M. charantia* leaf extracts.

In the current study, the primary antibody titer was found to be high in extract-treated group for 1000 mg/kg. This effect of enhancement of the antibody production by the extracts may be associated with effect on lymphoid cells as demonstrated by high mobilization of these cells. When the mice were sensitized with the bacteria, bacteria antigen was then taken up by macrophages and was processed. When a T lymphocyte sees the processed antigens on the B cell, the T cell then stimulates the B cells to undergo repeated cell divisions, enlargement and differentiation to form a clone of antibody secreted by plasma cells. Hence, the antibody then binds to the antigen, making them easier to ingest by the white blood cells. In the present study it has been demonstrated the secondary antibody has been increased 500 mg/kg and 1000 mg/kg treated-groups. This indicates enhanced responsiveness of macrophages, T and B lymphocytes involved in antibody synthesis by the extracts.

When *S. typhi* was given to mice, it has been found that the strain passes into blood and multiply. Using the extracts to avoid this blood infestation in mice, it was found the proportion of animal having bacterium in their blood decreases for certain concentration as it is for levamisole, a well-known immunostimulant drug. This experiment might be used as proof of the possible immunotherapeutic impact of *M. charantia* leaf extracts against *S. typhi* by simulating the phagocytic mechanisms or antibodies directed against such bacteria entering into the bloodstream, both of which have proven to be successful in eliminating or preventing blood from infection by microbes.

## Conclusion

The result of the current study suggests that immunomodulation may be a key factor in therapeutic activity of extract of *Momordica charantia* leaf in treatment of salmonellosis.

## Abbreviations

AcP: acid phosphatase
DMSO: dimethyl sulfoxide
FBS: fetal bovine serum,
LPS: lipopolysaccharide
MTT: 3- (4,5-dimetilthiazol-2-yl) −2.5-diphenyl tetrazolium bromide
NBT: nitroblue tetrazolium
NR: neutral red
OD: optical density
PBS: phosphate buffered saline
PM: peritoneal macrophages
*P*-NPP: para-nitrophenylphosphate
PN: Peritoneal neutrophils
RPMI: Roswell Park Memorial Institute
NO: Nitric oxide

## Declarations

*Ethics approval*

All animal handling protocols were performed following the guidelines in the Department of Biological Sciences, Faculty of Science, University of Bamenda, Cameroon which followed the « Principles of Laboratory Animal Care » from NIH publication Nos 85-23 approved by the ethic committee of the Cameroon Ministry of Scientific Research and Technology which has adopted the guidelines established by the European Union on Animal Care and Experimentation (CEE Council 86/609).

## Consent for publication

All authors have read and approved the publication of the manuscript.

## Availability of data and material

Data and material are available on needs.

## Author Contributions

**OM**: He performed the experiments, analyzed the data, wrote the paper, prepared figures and/or tables, and reviewed drafts of the paper, principal investigator. **HF**: She contributed analysis and experiments. **TC** and **KA**: They reviewed drafts of the paper.

## Acknowledgment

This research was done at the Institute of Agricultural Research for Development (IRAD) Bambui, Cameroon. The authors are grateful to the Director of IRAD and its collaborators. We are thankful to Director and its entire staff for his invaluable help in the conduction of this research work. We are thankful to Dr. Tacham W., Biological Sciences, Faculty of Science, University of Bamenda, for his invaluable help in the identification and collection of plant samples.

## References

1. Amirghofran Z. Herbal medicines for immunosuppression. Iran J Allergy Asthma Immunol, 2012, 11:111–119.

2. Alamgir M. and Uddin SJ. Recent advances on the ethnomedicinal plants as immunomodulatory agents. In: Chattopadhyay D, editor. Ethnomedicine: a source of complementary therapeutics. Fort P.O. Kerala, India: Research Signpost; 2010, 227–244.

3. Kumar V. and Sharma A. Neutrophils: Cinderella of innate immune system. Int Immunopharmacol., 2010, 10:1325–1334.

4. Eze JI and Ndukwe S. Effect of methanol extract of Mucuna pruriens seed on the immune response of mice. Comp Clin Pathol., 2012, 21:1343–1347.

5. Jantan I., Ahmad W, Bukhari S.N. Plant-derived immunomodulators: an insight on their preclinical evaluation and clinical trials. Front Plant Sci., 2015; 25; 6:655. 1 - 18.

6. Mahima AR, Rajib D, Latheef SK, Samad HA, Tiwari R, Vema AK, Dhama K. 2012. Immunomodulatory and therapeutic potentials of herbal, traditional/indigeneous and ethnoveterinary medicines. Pak J Biol Sci. 15:754–774.

7. Grover JK and Yadav SP. Pharmacological actions and potential uses of *Momordica charantia*: a review. Journal of Ethnopharmacology, 2004, 93 123–132

8. Shuo J., Mingyue S., Fan Z. and Jianhua X. Recent Advances in Momordica charantia: Functional Components and Biological Activities. Int. J. Mol.Sci., 2017, 18, 2555; 9–25.

9. Juvekar AR.; Hule AK.; Sakat SS.; Chaughule VA. In vitro and in vivo evaluation of immunomodulatory activity of methanol extract of Momordica charantia fruits. Drug Invent. Today, 2009, 1, 89–94.

10. Panda BC.; Mondal S.; Devi KSP.; Maiti TK; Khatua S.; Acharya K.; Islam SS. Pectic polysaccharide from the green fruits of Momordica charantia (Karela): Structural characterization and study of immunoenhancing and antioxidant properties. Carbohydr. Res., 2015, 401, 24–31.

11. Wang X.; Jin H.; Xu Z.; Gao L. Effects of momordica charantia L. saponins on immune system of senile mice. Acta Nutr. Sin., 2001, 263–266.

12. Deng YY; Yi Y; Zhang LF.; Zhang RF.; Zhang Y.; Wei ZC.; Zhang MW. Immunomodulatory activity and partial characterization of polysaccharides from Momordica charantia. Molecules, 2014, 19, 13432–13447.

13. Liu JQ; Chen JC.; Wang CF; Qiu MH. New cucurbitane triterpenoids and steroidal glycoside from Momordica charantia. Molecules 2009, 14, 4804–4813.

14. Najafi P.; Torki M. Performance, blood metabolites and immunocompetaence of broiler chicks fed diets included essential oils of medicinal herbs. J. Anim. Vet. Adv. 2010, 9, 1164–1168.

15. Ayeni MJ; Oyeyemi SD; Kayode J; Peter GP. Phytochemical, proximate and mineral analyses of the leaves of Gossypium hirsutum L. and Momordica charantia L. J. Nat. Sci. Res. 2015, 5, 99–107.

16. Chen J.; Tian R.; Qiu M.; Lu L.; Zheng Y.; Zhang Z. Trinorcucurbitane and cucurbitane triterpenoids from the roots of Momordica charantia. Phytochemistry. 2008, 69, 1043–1048.

17. Zhao GT; Liu JQ; Deng YY; Li HZ; Chen JC; Zhang ZR.; Qiu MH. Cucurbitane-type triterpenoids from the stems and leaves of Momordica charantia. Fitoterapia, 2014, 95, 75–82.

18. Chang CI; Chen CR; Liao YW; Cheng HL; Chen YC; Chou CH. Cucurbitane-type triterpenoids from the stems of Momordica charantia. J. Nat. Prod. 2008, 71, 1327–1330.

19. Begum S.; Ahmed M.; Siddiqui BS; Khan A.; Saify ZS.; Arif M. Triterpenes, a sterol and a monocyclic alcohol from Momordica charantia. Phytochemistry, 1997, 44, 1313–1320.

20. Akihisa T.; Higo N.; Tokuda H.; Ukiya M.; Akazawa H.; Tochigi Y.; Nishino H. Cucurbitane-type triterpenoids from the fruits of Momordica charantia and their cancer chemopreventive effects. J. Nat. Prod. 2007, 70, 1233–1239.

21. Ma L.; Yu AH; Sun LL; Gao W.; Zhang MM; Su YL; Liu H; Ji T. Two new bidesmoside triterpenoid saponins from the seeds of Momordica charantia L. Molecules, 2014, 19, 2238–2246.

22. Murakami T.; Emoto A.; Matsuda H.; Yoshikawa M. Medicinal foodstuffs. XXI. Structures of new cucurbitane-type triterpene glycosides, goyaglycosides-a,-b,-c,-d,-e,-f,-g, and-h, and new oleanane-type triterpene saponins, goyasaponins I, II, and III, from the fresh fruit of Japanese Momordica charantia L. Chem. Pharm. Bull., 2001, 49, 54–63.

23. Ahmad Z; Zamhuri, KF; Yaacob A; Siong CH; Selvarajah M; Ismail A.; Hakim MN. In vitro anti-diabetic activities and chemical analysis of polypeptide-k and oil isolated from seeds of Momordica charantia (bitter gourd). Molecules, 2012, 17, 9631–9640.

24. Wen LJ.; Liu WF. Study on extracting and antioxidant activity of flavonoids from Momordica charantia L. Food Sci. 2007, 9, 042.

25. Okabe H.; Miyahara Y.; Yamauchi T; Miyahara K.; Kawasaki T. Studies on the constituents of Momordica charantia LI Isolation and characterization of momordicosides A and B, glycosides of a pentahydroxy-cucurbitane triterpene. Chem. Pharm. Bull. 1980, 28, 2753–2762.

26. NIH. 1996. Guide for the care and use of laboratory animals. Washington, DC: National Academy Press.

27. Aurasorn S., Pattana S. Anti-inflammatory activity of a Vernonia cinerea methanolic extract in vitro. ScienceAsia, 2015, 41: 392–399.

28. Margot L., Urban J-LL., Per G., and Per C. Human Alveolar Macrophage Phagocytic Function is Impaired by Aggregates of Ultrafine Carbon Particles. Environmental Research Section A, 2001, 86:244 – 253.

29. Oumar M., Tume C., Ateufack G., Ngo TG and Kamanyi A. Immunological In Vivo and In Vitro Investigations of Aqueous Extract of Stem Bark of Pterocarpus erinaceus Poir (Fabaceae). Am J Med Sci., 2018; 356 (1): 56 – 63.

30. Ribeiro RA, Flores CA, Cunha F, Ferreira S. 1991. IL-8 causes in vivo neutrophil migration by a cell-dependent mechanism. Immunology. 73:472–477.

31. Eun-Jin Y., Eun-Young Y., Gwanpil S., Gi-Ok K., Chang-Gu H. Inhibition of nitric oxide production in lipopolysaccharide-activated RAW 264.7 macrophages by Jeju plant extracts. Interdisc Toxicol. 2009; Vol. 2(4): 245–249.

32. Adams L., Franco MC and AG. Estevez. Reactive nitrogen species in cellular signaling. Exp. Biol. Med. (Maywood)., 2015; 240:711–717.

33. Förster R, Sozzani S. 2013. Emerging aspects of leukocyte migration. Eur J Immunol., 43:1404–1406.

